# "Smurf Mice": revolutionising our understanding of age-related and end-of-life animal physiology

**DOI:** 10.1101/2025.06.14.659669

**Authors:** Céline Cansell, Vivien Goepp, Fanny Bain, Nicolas Todd, Veronique Douard, Magali Monnoye, Flaminia Zane, Clara Sanchez, Nicolas Pietrancosta, Carole Rovere, Raphaël GP Denis, Serge Luquet, Michael Rera

## Abstract

Living animals reach their end-of-life through a stereotypic set of fascinating but poorly understood processes. The discovery, first in flies and later in nematodes and zebrafish, of the "Smurf phenotype" is a central tool for picking this complex "lock of biology", that one of ageing. Using the Smurfs, we have shown an evolutionarily conserved end-of-life transition across Drosophilids, nematodes and zebrafish. This tool has been key to identify the discontinuous nature of ageing and predict impending death from natural causes as well as from environmental stresses. This phenotype allowed us to discover that ageing is made up of two successive phases : a first phase where individuals are healthy and have no risk of mortality, but show an age-dependent and increasing risk of entering a second phase, characterized by the so-called hallmarks of ageing and a high risk of death. Here, we test whether these two consecutive phases of ageing separated by the Smurf transition are a conserved feature of ageing in the mammals using *Mus musculus as a* model. We performed a longitudinal longevity study using both males and females from two different mouse genetic backgrounds and by integrating physiological, metabolic and molecular measurements with the life history of approximately 150 mice. We show the existence of a phenotypic signature typical of the last phase of life, observable at any chronological age. Validating the two-phase ageing model in a mammalian organism allows better characterized the high risk of imminent death and would extend its implications to a broader range of species for ageing research.

## Introduction

In 2015, building upon the first description of a pre-death phenotype we termed Smurf^1^, we further validated its relevance by developing a new mathematical model for analyzing longevity data. This model of ageing, first described in Drosophila, is characterized by - at least - two successive and necessary phases, a phase 1 where individuals are healthy and have no risk of mortality but an age-dependent increasing risk of entering phase 2, and a phase 2 where individuals show so-called hallmarks of ageing^2,3^ as well as a high risk of death. The discovery of this model in Drosophila was made possible by a singular characteristic of phase 2, its increased intestinal permeability^1^. To assess intestinal permeability in Drosophila, a non-absorbed and non-toxic blue food dye (FD&C blue dye #1) is added to the medium fed to adult individuals. In phase 1, this dye cannot pass the intestinal barrier and remains in the digestive tract. In contrast, for individuals in phase 2, the dye crosses the intestinal epithelium by mechanisms that are still unknown and reaches the fly circulation. The blue color is visible to the naked eye and blue individuals were hence named "Smurfs"^3^. Individual flies of both sexes monitored daily showed that about 50%^4^ to 100%^1,5^ of individuals go through these two consecutive phases^1,6^. These include decreased energy storage (triglycerides, glycogen)^1^, locomotor activity^1^ and fertility^7^, a deregulation of insulin signaling^3^, and an increase in systemic inflammation^1^. This powerful predictor - the Smurf phenotype - of an individual’s death strongly supports that instead of describing ageing as a progressive and continuous age-related alteration of the physiology in all individuals within a population, we should rather view it as an abrupt alteration in an age-dependent growing proportion of individuals with high mortality risk. We hereby propose to use the Smurf phenotype as an objective means to identify frail individuals.

The Smurf two-phase ageing model groups several important ideas with respect to ageing: (1) not all individuals of the same chronological age have the same physiological age, (2) although some parameters may evolve gradually, the transition from phase 1 to phase 2 is abrupt, and (3) it is possible to identify physiologically old individuals, i.e. at higher risk of impending death than the rest of the population., This model of ageing is evolutionarily conserved as it was also described in nematodes^8,9^ (*Caenorhabditis elegans*) and fish^8^ (*Danio rerio*). Additionally to its ability to predict a high-risk of impending natural death in Drosophila, Smurf phenotype was used to show death prediction in non-naturally occurring causes of death such as a traumatic brain injury Drosophila model^10^ but also acute oxidative stress^1^. Studies regarding age-dependent increase in intestinal permeability in aged rodents^11–13^, aged primates^14^, and in critically ill humans in Intensive Care Unit^15,16^ led us to investigate whether this “Smurf’’ two-phase ageing model is also conserved in mammals. Moreover, in humans, certain phenotypic markers such as physical capability^17^, sense of smell^18^ and facial appearance^19^ have been described as predictors of death. Recently, studies have identified metabolic^20^, protein^21^ and inflammatory biomarkers^22^ that can identify, within a population of the same chronological age, individuals with a high risk of mortality from any causes. This opens up the exciting possibility that the same biological features preceding death could be really conserved between flies and humans.

In the present study, we decided to assess the conservation of our predicting model in *Mus musculus*. We thus looked for a phenotypic signature that would be associated with increased intestinal permeability prior to death, and observable at any chronological age. For this, we use two mouse lines, with different lifespans: the AKRJ line with a T_50_ (the age at which 50% of a population has died) of 9 months and the C57Bl6/J line with a T_50_ of 30 months^23^. We regularly assess intestinal permeability, metabolic parameters, inflammatory status and microbiota homeostasis throughout the life of each individual until their natural death. This approach allows us to answer the following four research questions: (1) Is the end of life in mice characterized by an increase in intestinal permeability? (2) Is there a specific physiological signature of the end of life in mice, independent of the chronological age(3) Do defined biomarkers allow us to discriminate between two subpopulations in mice characterized by different mortality risk at any chronological age? (4) How well can we predict impending death in mice?

The demonstration that the two-phase ageing model initially described in flies also applies in mice would represent multiple benefits for the ageing research field. First, because it would prove the two-phase model of ageing to be a “public” path of ageing and provide arguments to pursue this research in humans. Secondly, it would make possible the assessment of an individual’s physiological age using a method validated across multiple model organisms. Third, the multiple physiological parameters that will be followed longitudinally until the natural death of individuals, will allow the construction of a new multiparametric predictive model^24^, based on metabolic criteria likely to be relevant across model organisms. Finally, it would also allow a better understanding of the underlying mechanisms of ageing by offering the possibility to study the onset of the aforementioned hallmarks of ageing in healthy, phase 1 individuals.

## Results

### Intestinal permeability is increased towards mice end-of-life

In Drosophila, phase 2 of ageing is characterized by an increased intestinal permeability, which can be easily detected by the leakage of a blue food dye through the intestinal epithelium into the hemolymph, resulting in a blue coloration of the flies^1^. To explore whether a similar phenomenon occurs at the end of life in mice, we studied three cohorts: AKR/J females, representing pathological ageing, and C57BL/6J males and females, representing normal ageing. AKR/J mice have a median lifespan (T_50_) of approximately 9 months^23^, while C57BL/6J mice have a T_50_ of about 30 months^23^ (Supplementary Figures 1A, 1B and 1C) The shorter lifespan of AKR/J mice is due to congenital viremia, which consistently progresses to leukemia or lymphoma^25^. Previous studies in Drosophila suggest that factors leading to reduced lifespan, whether pathological or environmental, do not prevent the appearance of the Smurf phenotype but rather accelerate its onset^1,5^.

All mice were aged under standard conditions and tested every 2 to 3 weeks for intestinal permeability using the FITC-Dextran gavage assay^26^. When the results were plotted against chronological age, an increase in intestinal permeability was observed at older ages in the AKR/J female cohort (Figure 1Ai), whereas no clear age-related changes in permeability were detected in either of the two C57BL/6J cohorts (Figure 1Bi and Supplementary Figure 1Di). To investigate potential shifts occurring shortly before death, we re-analyzed the data using the R package “segmented” to fit a Segmented Linear Model (SLM). This analytical approach has not been applied for longitudinal physiological parameters before. The analysis followed a stepwise procedure: a linear model was first fitted and its adjusted R² calculated; this was followed by a comparison between models with zero and two breakpoints. If the test was significant (p < 0.05), an additional comparison between models with one and two breakpoints was performed. Finally, the segmented model was fitted, and its adjusted R² compared with that of the linear model. A higher adjusted R² for the segmented model indicated a change in data behavior, with a breakpoint marking a shift in intestinal permeability before death. This analysis identified a breakpoint in each group: 31 days before death in AKR/J females (Figure 1Aii), 17 days before death in C57BL/6J females (Figure 1Bii), and 10 days before death in C57BL/6J males (Supplementary Figure 1Dii).

**Figure 1:**
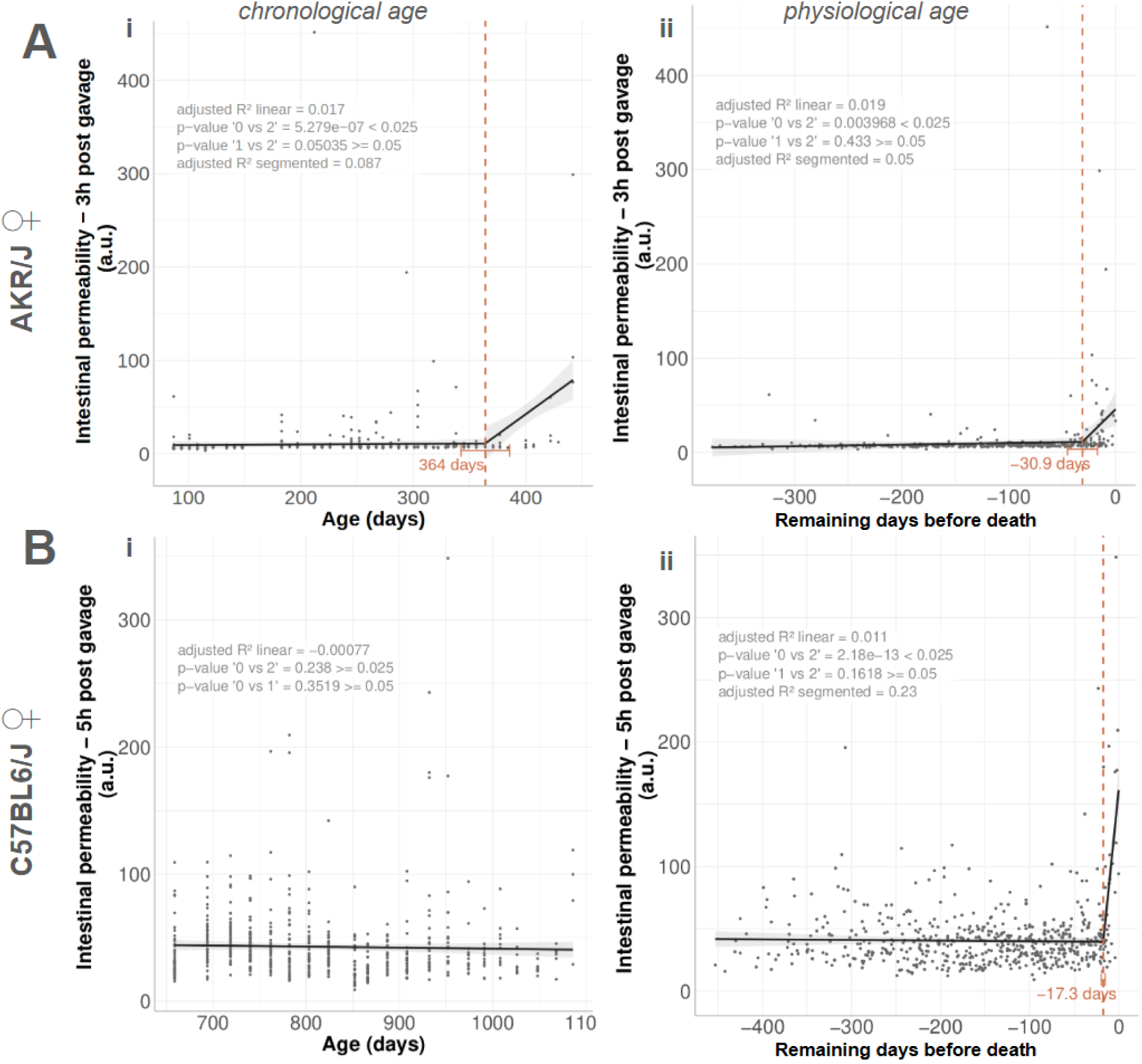
Longitudinal assessment of intestinal permeability until natural death in C57BL/6J and AKR/J female mice cohorts. Fluorescence unit in plasma 3h after FITC-dextran gavage indicating intestinal permeability in a cohort AKR/J females (n=45) and (B) C57BL6/J females (n=48) as a function of chronological age of individuals (i) and as a function of physiological age (ii) meaning days remaining before natural death of individuals. The dataset was modified to alleviate the impact of inter-individual variability by negating the intercept, slope, and slope difference. This treatment does not affect the pre-death pattern of the data. The vertical dashed orange line shows the breakpoint identified by segmented linear model (SLM)^27^ analysis of longitudinal intestinal permeability values. Black lines represent the results of SLM analysis before and after the breakpoint. Results of subsequent calculations and statistical tests (p-values) are written on the graph : adjusted R^2 for the linear and segmented fit ; statistical test for the absence of 1 breakpoint (BP) and for the absence of a 2nd BP (Pseudo-score test).

These results demonstrate that, as in Drosophila, a late-life increase in intestinal permeability occurs in mice during the final weeks before death.

### An abrupt physiological shift marks the end of life in mice

To determine whether a physiological signature precedes death in mice, we conducted longitudinal analyses of several parameters, including - we mention the parameters that are later used for modeling - glycemia, body weight, food intake, fat mass, and body temperature, in the three cohorts : AKR/J and C57BL/6J females (Figures 2 and 3) as well as C57BL/6J males (Supplementary Figure 2). We plotted data for each variable against chronological age (Figures 2Ai and 2Bi) and physiological age, defined as time to death (Figure 3 and Supplementary Figure 2), and applied segmented linear modeling (SLM) to identify potential breakpoints in the trajectories.

**Figure 2:**
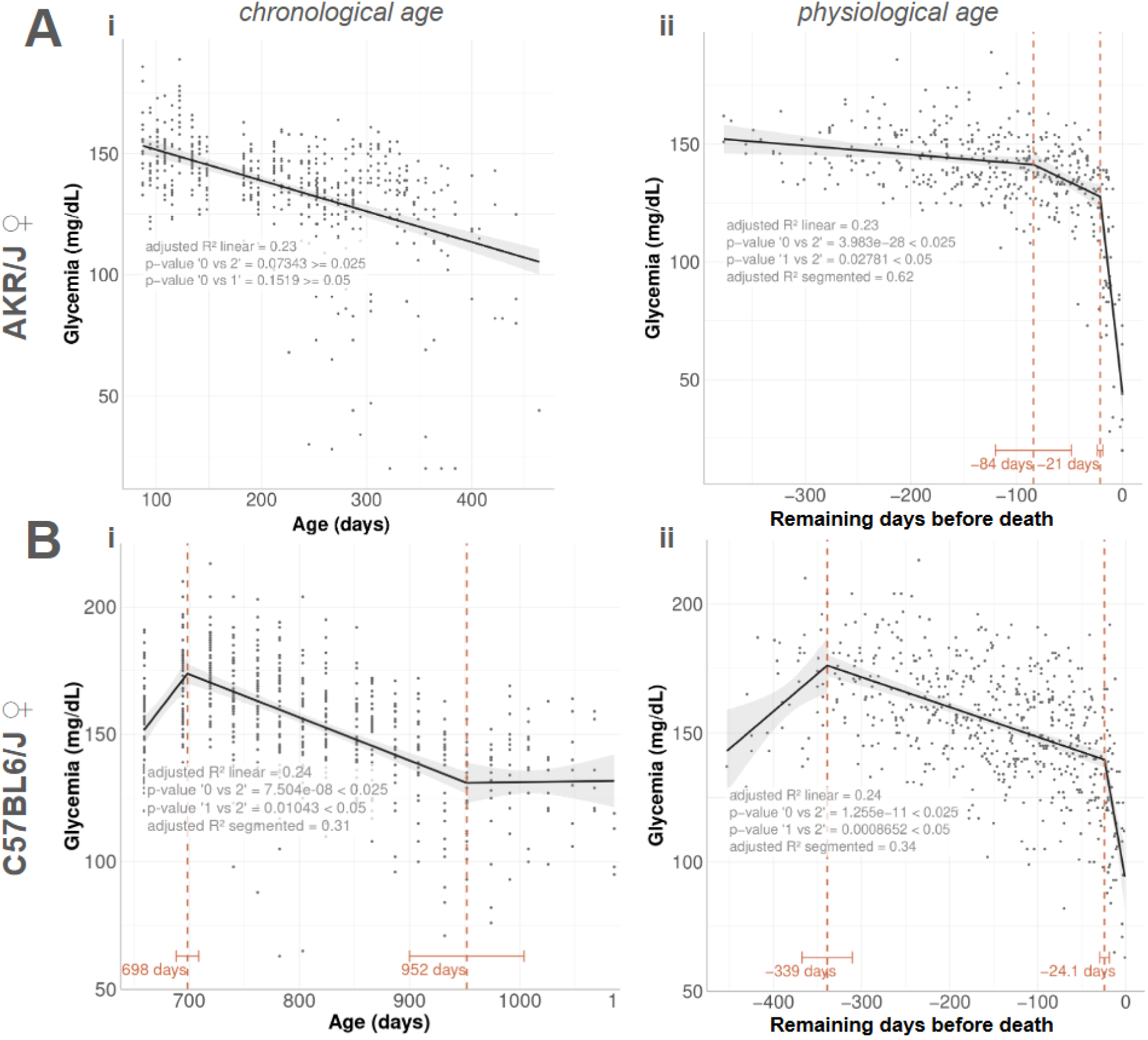
A clear and sharp drop in glucose levels occurs shortly before death in C57BL/6J and AKR/J female mice cohorts. Longitudinal representation of glucose levels in plasma in a cohort (A) AKR/J females (n=45) and (B) C57BL6/J females (n=48) as a function of chronological age of individuals (i) and as a function of physiological age (ii) meaning days remaining before natural death of individuals. The dataset was modified to alleviate the impact of inter-individual variability by negating the intercept, slope, and slope differences. This treatment does not affect the pre-death pattern of the data. The vertical dashed orange line shows the breakpoint identified by segmented linear model (SLM) analysis of longitudinal intestinal permeability values. Black lines represent the results of SLM analysis before and after the breakpoint. Results of subsequent calculations and statistical tests (p-values) are included on the graph: adjusted R^2 for the linear and segmented fit ; statistical test for the absence of 1 breakpoint (BP)^28^ and the absence of a 2nd BP (Pseudo-score test).

**Figure 3:**
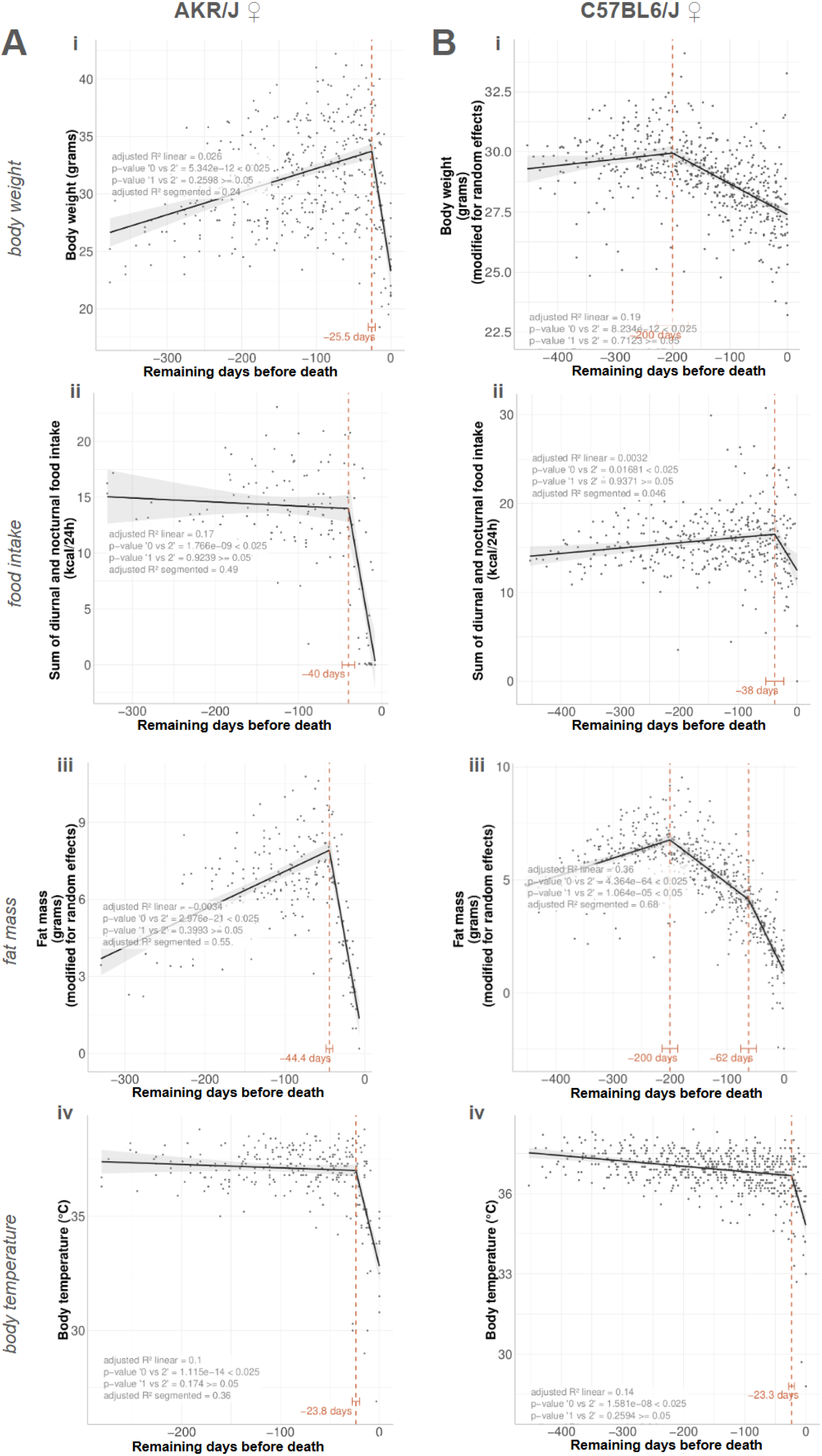
Multiple physiological parameters undergo end-of-life–specific deterioration in female C57BL/6J and AKR/J mouse cohorts. Segmented linear model analysis of physiological variables in a cohort (A) AKR/J females (n=45) and (B) C57BL6/J females (n=48). Longitudinal representation of body weight (i), food intake (ii), fat mass (iii) and rectal temperature (iv) all expressed as a function of time before recorded death. The dataset was modified to alleviate the impact of inter-individual variability by negating the intercept, slope, and slope differences. This treatment does not affect the pre-death pattern of the data. The vertical dashed orange line shows the breakpoint identified by segmented linear model (SLM) analysis of longitudinal intestinal permeability values. Black lines represent the results of SLM analysis before and after the breakpoint. Results of subsequent calculations and statistical tests (p-values) are written on the graph : adjusted R^2 for the linear and segmented fit ; statistical test for the absence of 1 breakpoint^28^ and the absence of a 2nd BP (Pseudo-score test).

When we analysed glycemia across chronological age, no breakpoint was detected in AKR/J females (Figure 2Ai). In C57BL/6J females, breakpoints were identified, reflecting an initial decrease followed by a plateau (Figure 2Bi). Those trajectories may suggest a period of stability in the final months of life; however, replotting the data against physiological age revealed a different pattern. In this representation, glycemia remained relatively stable until a distinct and sharper decline occurred shortly before death, around 21 days in AKR/J females and 24 days in C57BL/6J females (Figure 2Aii and 2Bii). These changes were supported by a significantly better fit of the segmented model compared to a linear model, indicating a marked shift in glycemia near the end of life.

Analysis of additional physiological parameters showed comparable trends in the weeks preceding death (Figure 3). In AKR/J females, significant breakpoints were detected for body weight (−25 days), food intake (−40 days), fat mass (−44 days), and body temperature (−24 days). In C57BL/6J females, breakpoints were observed for food intake (−38 days) and body temperature (−23 days). A reduction in body weight and fat mass was also seen in C57BL/6J females prior to death, although the decline was more gradual and the abrupt drop observed in AKR/J mice was less pronounced. A similar pattern of physiological changes was observed in C57BL/6J males (Supplementary Figure 2).

Together, these results show that multiple physiological variables undergo synchronized changes during the final phase of life. This supports the existence of a reproducible and conserved physiological transition preceding death in mice. For the remainder of this article, we will refer to two distinct phases of ageing: ’phase 1’ before the breakpoint and ’phase 2’ after it.

### Biomarkers identified through a “near-death” physiological state stratifies individuals by mortality risk

We identified two distinct and consecutive phases—Phase 1 and Phase 2—during the ageing process in mice, based on the biphasic behavior of several physiological parameters. This enabled us to investigate whether these parameters could be used to determine an individual’s ageing phase independently of chronological age. To this end, we selected six key variables that display a biphasic trajectory and are consistently observed across female mouse cohorts: intestinal permeability, body weight and temperature, food intake, total fat mass, and glycemia.

We followed the workflow presented in Figure 4A. First, to identify the transition point between phases 1 and 2 of ageing, we calculated the mean breakpoint (BP) for these five key variables, determined using the segmented linear regression model (SLM) described earlier. Each cohort was thus assigned an average BP (AKR/J females = 31 days; C57BL6/J females = 60 days; C57BL6/J males = 31 days). Subsequently, we performed a principal component analysis (PCA) with these five variables of interest, and used the score from the first dimension (Dim1) to create a composite score encompassing these physiological parameters for a given chronological age. To categorize individuals as non-Smurf (phase 1 of ageing) or Smurf (phase 2 of ageing), we established a threshold value for Dim1, using the mean value of this score during phase 1 of ageing as a reference (see threshold calculation step in Figure 4A). This categorization allowed us to demonstrate that, for all three cohorts of mice, the survival time of individuals categorized as Smurf (AKR/J females = 29 days ± 5 days; C57BL6/J females = 32 days ± 5 days; C57BL6/J males = 53 days ± 13 days; Figures 4Bii and 4Cii; Supplementary Figure 3Aii), regardless of their chronological age, is significantly shorter than their life expectancy at birth (AKR/J females = 10 months ± 12 days; C57BL6/J females = 30 months ± 17 days; C57BL6/J males = 30 months ± 16 days; Figures 4Bi and 4Ci; Supplementary Figure 3Ai). As previously described in Drosophila, the life expectancy of Smurf individuals is significantly shorter than that of non-Smurf individuals of the same chronological age (Figures 4Biii and 4Ciii; Supplementary Figure 3iii). Furthermore, the proportion of Smurf individuals increases with chronological age (Figures 4Biv and 4Civ; Supplementary Figure 3Aiv). Although the time-dependent increase of the Smurf proportion was previously described as approximately linear in nematodes, Drosophila and zebrafish, it is significantly better fitted by an exponential in mice.

**Figure 4:**
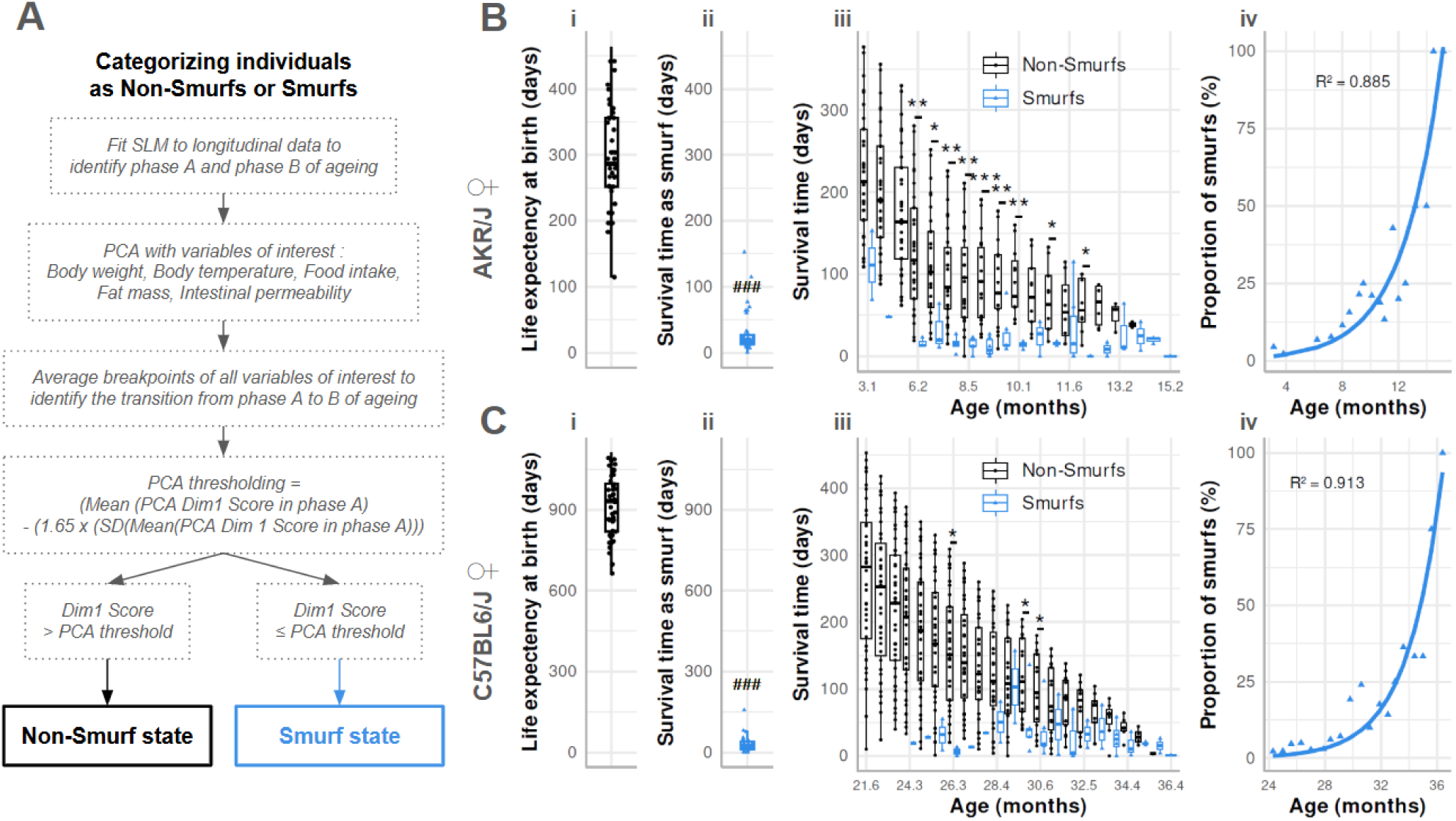
Identifying differential mortality risk in female C57BL/6J and AKR/J mice of the same chronological age. (A) Graphical representation of logical events used to classify individuals as non-smurf or smurf (see “Materials & Methods” for details). (B) & (C) (i) Individual life expectancy at birth in days (black boxes, points); (ii) Individual survival time as smurf in days (blue boxes, triangles); (iii) Individual survival time as non-smurf (black boxes, points) and as smurf (blue boxes, triangles) as a function of chronological age; (iv) Proportion of smurf individuals as a percentage of the living population (blue triangles), with an exponential growth regression of smurf proportion (blue line). Data are presented for AKR/J female mice (n=45) (B) and C57BL/6J female mice (n=48) (C). The box plot represents the interquartile range, showing the middle 50% of the data from the first to the third quartile. The line inside the box marks the median, and whiskers extend to the smallest and largest values. Statistical significance: ### p < 0.001 by paired Wilcoxon test comparing mean life expectancy at birth and mean survival time as smurf (ii); *p < 0.05, **p < 0.01, ***p < 0.001 by Wilcoxon test comparing mean survival times of non-smurf and smurf individuals (iii). In the AKR/J female cohort, 7 individuals were never classified as smurfs, and 2 out of 38 showed a reversion to the non-smurf state after their initial categorization as smurf. In the C57BL6/J female cohort, 24 individuals were never classified as smurfs, and 5 out of 24 showed a reversion to the non-smurf state after their initial categorization as smurf. SLM = Segmented Linear Model; PCA = Principal Component Analysis; Dim1 = Dimension 1 of the PCA ; R2 = 1 - (sum squared regression / total sum of squared).

We then represented mice followed up from adulthood to their natural death as individual life histories centered on the moment of the Smurf transition (Supplementary Figure 3B, 3C, 3D). This representation helps visualizing that (i) individuals become Smurf at different, unpredictable, times, (ii) there is no correlation between the age at smurf transition and the remaining life expectancy of a smurf and (iii) the remaining lifespan of individuals in Phase 2 of ageing (after Smurf transition) is significantly decreased compared to that of Phase 1 of ageing (before Smurf transition).

All these findings are consistent across AKR/J females (Figure 4B), C57BL6/J females (Figure 4C) and males (Supplementary Figure 3). Taken together, these results demonstrate that it is possible to distinguish two subpopulations of mice, each characterized by a different mortality risk, regardless of chronological age, on the basis of the combined levels of the six biomarkers we have identified above.

### Smurfness-associated biomarker predicts impending death in mice

Based on Smurfness-associated biomarker, we build a predictive model for high risk of impending death. To do so, we developed a joint model of biomarker trajectories and survival^29^ (see the ’Joint Model’ subsection under ’Statistical Analysis’ in Materials and Methods). Included biomarkers are the ones that consistently displayed a breakpoint in their values before death across the three mouse cohorts (AKR/J females, C57BL/6J females, and C57BL/6J males). These variables are: body temperature, fat mass, glycemia and intestinal permeability. Results of the joint models are presented in Table 1. When the coefficient is estimated to be positive, a higher biomarker value is associated with a higher force of mortality. Aside from intestinal permeability, we expect a decline in biomarker value to be associated with higher mortality, we expect negative coefficients. Note that p-values reported in Table 1, defined as 2min(P(θ<0|y), P(θ>0|y)) for each coefficient θ, are two-tailed. Thus, the posterior probability that the effect of, e.g., body temperature is <0 is 1-pvalue/2: a p-value of 0.06 corresponds to a 1-0.06/2=0.97 probability of a negative effect of body temperature (i.e. lower body temperature associated with a higher risk of death). In both C57BL6 and AKRJ mice, low glycemia was found to be associated with an increased risk of death; the estimated effect was found to be close in both strains. We achieved a higher precision of the estimation in C57 mice thanks to a larger sample size. In C57 mice, evidence was also found that higher intestinal permeability contributed a higher risk of death (p=0.02) ; some evidence was also found for an effect of low body temperature on higher mortality (p=0.06).

**Table 1:**
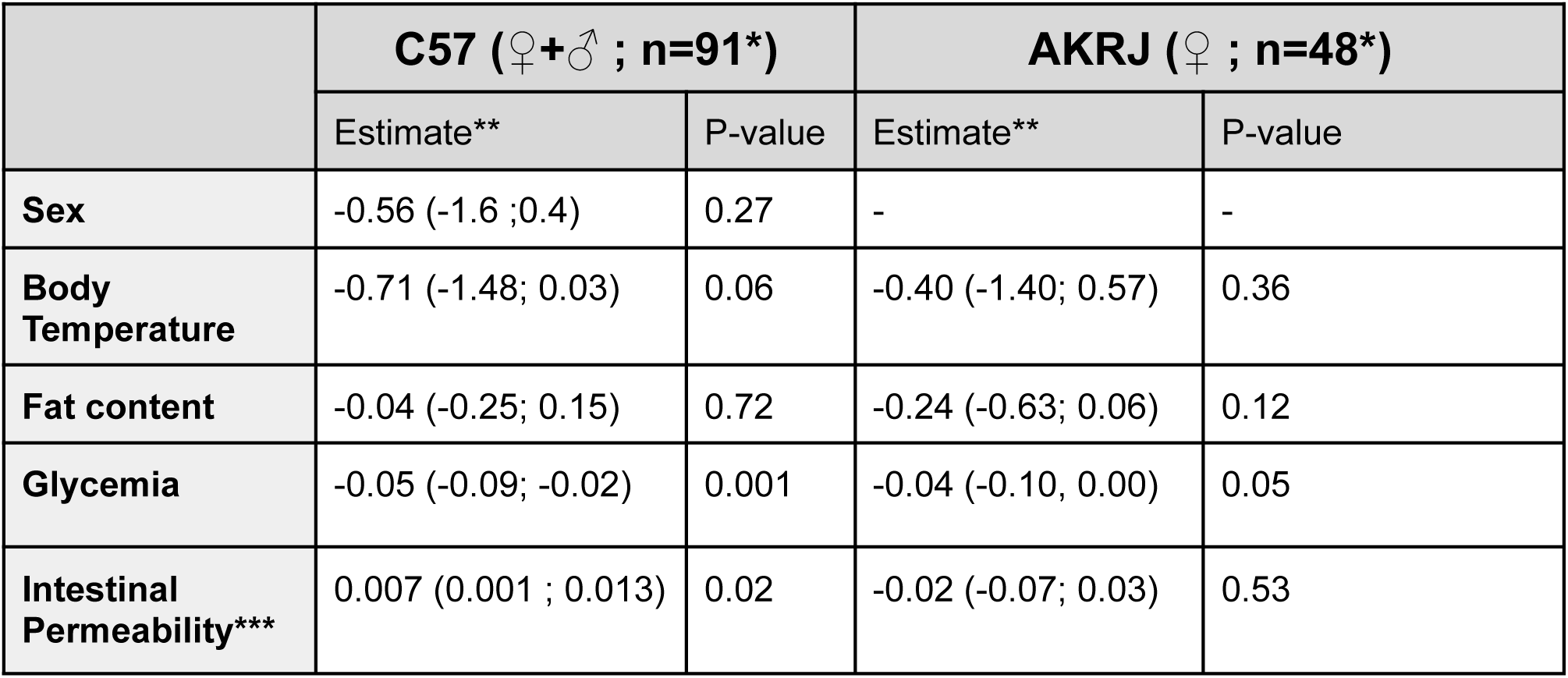
Effect of covariates on the force of mortality. For the four biomarkers, current value is entered in the predictor of the force of mortality (the instantaneous risk of death). *To build the model, each mouse required at least one value for each variable, which is why C57BL/6J male 42 and AKR/J females 27 and 50 were excluded.**mean posterior (95% Uncertainty Interval). *** C57: 5 hour intestinal permeability; AKRJ: 3 hour intestinal permeability.

For the purpose of illustration, we also show on Figure 5 dynamic predictions for two selected mice from the C57BL/6J female cohort, mice 1 and 14. Those two mice were chosen among C57BL6/J females still alive at age 1 000 days, based on a visual examination of their biomarker profiles up to age 1 000 days only : mouse 14 as an apparently healthy individual, mouse 1 as an individual with a profile of deteriorating biomarkers. Prediction of remaining lifespan was based only on biomarker profiles up to age 1 000 days. As shown on Figure 5, the joint model predicted significantly shorter lifespan for mouse 1. Death indeed occurred at age 1010 for mouse 1, while mouse 14 outlived it by approximately 3 months.

**Figure 5:**
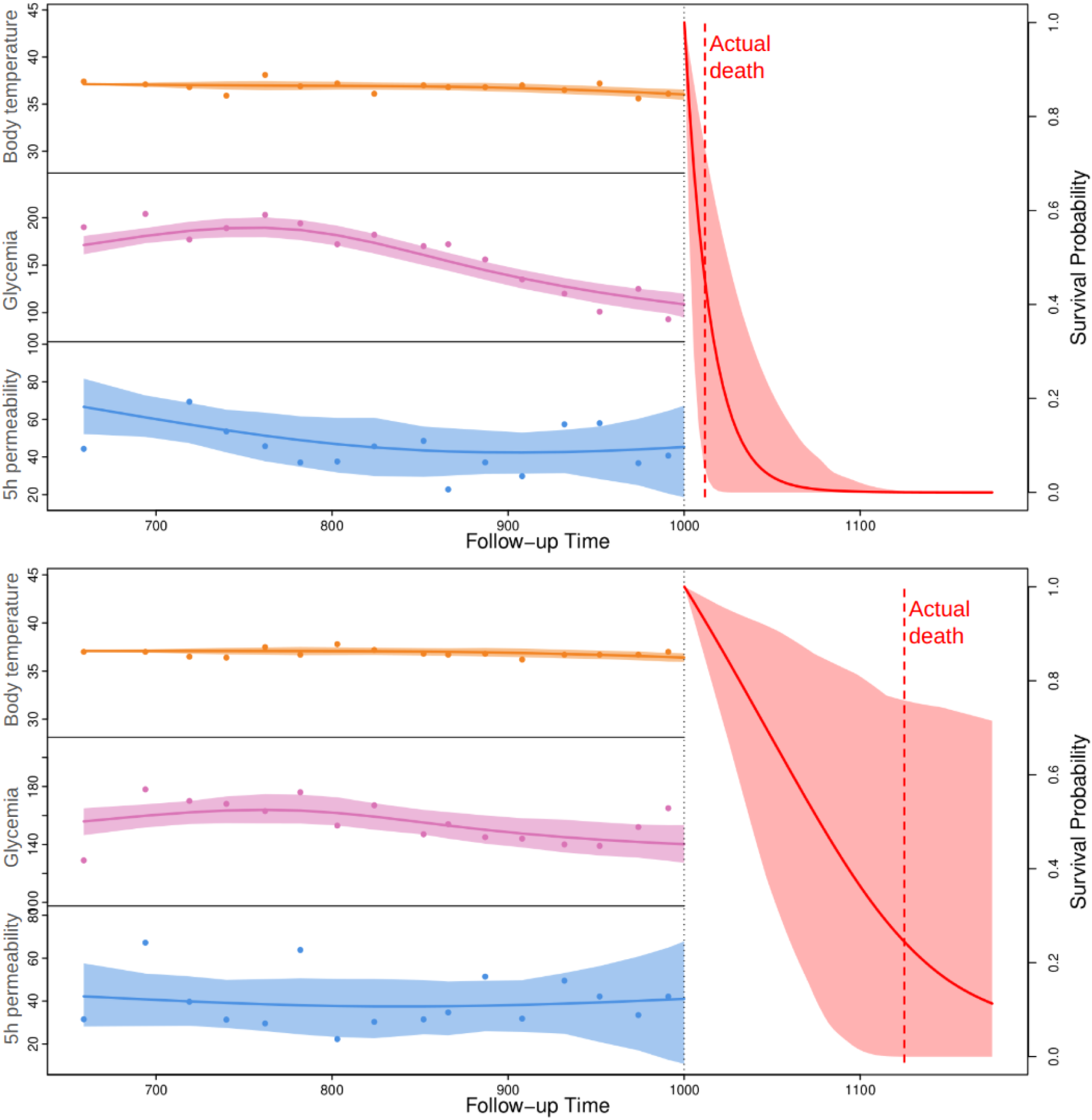
Predicting risk of impending death in 1000 day-old C57BL6/J females. Top: individual with a profile of apparently deteriorating biomarkers. Bottom: apparently healthy individual based on biomarker profile. Biomarker profiles up to age 1 000 days are shown (left). The prediction is given as an estimated survival function from age 1 000 days onwards, with 95% Uncertainty Interval (right).

Taken together with the results presented along the present article, this last modeling attempt suggests that we are able, with only three of the biomarker followed in our study, to discriminate individuals about to die from natural causes from their siblings across different strains.

## Discussion

In 2011, Rera et al. first showed that *in vivo* intestinal permeability to a non-toxic, non-absorbed blue food dye (FD&C blue #1) is a reliable marker of physiological age in *Drosophila melanogaster*^30^. This marker proved more relevant than chronological age in predicting several ageing-related phenotypic traits, such as loss of energy reserves, systemic inflammation, reduced motility, and increased mortality^1^. The first phase is a silent period that covers most of the organism’s life and during which the age-related risk to become Smurf increases. The second phase, referred to as the Smurf phase due to the blue coloration observed in flies, begins abruptly and is defined by the highly correlated onset of multiple markers of advanced ageing^30^. This transition, independently of the chronological age at which it occurs, is highly predictive of impending death. Based on these observations, we proposed a two-phase ageing model known as the 2PAC model^5^. More recently, we showed that Smurf flies are the only ones to show the transcriptional hallmarks of ageing to be significantly altered, independently of their chronological age^6^. We considered that validating the existence of this two-phase process in a mammalian model would be relevant for advancing research in the field of ageing.

Our initial hypothesis was that the two-phase ageing model originally described in Drosophila would be conserved in mice. To test this hypothesis, we addressed and provided evidence for the following points: (1) the end of life in mice is characterized by an increase in intestinal permeability; (2) a specific physiological transition precedes death; (3) defined biomarkers can discriminate between two subpopulations of mice with different mortality risks, regardless of chronological age; and (4) it is possible to accurately predict imminent death in mice.

We observed that in both male and female C57BL/6J mice, as well as in female AKR/J mice, ageing could be divided into two consecutive phases with distinct physiological features. Their last 30 days of life include notably, an increased intestinal permeability (Figure 1 and Supplementary Figure 1), a drop in glycemia, reduced food intake (not in males), and impaired body temperature regulation. We previously described sexual dimorphism that is species-specific^1,8^ in the Smurf phenotype, without yet being able to propose an explanation. Those changes occurred within a timeframe of two to three weeks and concurrently, with the likelihood of these changes increasing over time (Figure 2—3 and Supplementary Figure 2). In AKR/J mice, the “Smurf package”, that is the whole set of phenotypes associated with that end-of-life phase, was associated with a marked reduction in body weight primarily driven by fat mass depletion (Figure 3), which may account for the pronounced decrease in plasma leptin levels observed in the Smurf group (Supplementary Figure 4). A similar phenomenon was observed in C57BL/6J female mice, although it tended to occur slightly earlier relative to the other physiological parameters and in a less abrupt manner (Figure 3). This loss of fat storage is reminiscent of what was first described in Smurf flies^1^. In all cases, individuals undergoing this transition exhibited the typical characteristics of aged individuals^2^, indicating disrupted metabolism and a high risk of imminent death, with remaining lifespan largely independent of chronological age at the time of transition (Figure 4 and Supplementary Figure 3). Smurf mice also exhibit a classical feature of aging^2^ - chronic low-grade inflammation - characterized by a significant increase in circulating inflammatory markers such as IL-10, RANTES, and TNF-α compared to non-Smurf individuals (Supplementary Figure 4). Using a joint model approach, we built a mortality prediction model that we tested on mice above age 1000 and observed that low blood glucose levels are strongly associated with an increased risk of death in mice (Table 1). These results corroborate and extend the study by Palliyaguru et al.^31^, who, through a mortality analysis based on metabolic stratification, demonstrated that reduced plasma glucose levels are linked to higher mortality in mouse models. The observed hypoglycemia in our cohort may be partially attributed to reduced food intake (Figure 3), but also likely reflects broader disruptions in hormonal regulation responsible for maintaining glucose homeostasis, as suggested by the decrease in plasma insulin levels observed in the AKR/J cohort (Supplementary Figure 4).

Some physiological changes appeared specific to sex or genetic background. However, the “Smurf package”, when detected, consistently appeared suddenly across all studied organisms. Our inability to identify pre-Smurf individuals limited our capacity to detect the early events of the transition, whose duration appears to scale with the species’ life expectancy^6^. An important observation is that these two ageing phases occurred both in the context of “normal” ageing (as observed in the C57BL/6J cohorts) and in a pathological context (as observed in AKR/J mice). This aligns with our previous results, as well as those of others, showing that the Smurf phase constitutes a stereotyped path to death under various environmental conditions^1,32,33^, genetic alterations^1,5,30,34^, or physical trauma^10^.

A major limitation of the study is that not all individuals in our study underwent a detectable Smurf phase before death. Moreover, increased intestinal permeability immediately preceding death was not observed in all individuals. One limitation that may account for this is the relatively low frequency of intestinal permeability assessment - once every 2 to 3 weeks - which roughly matches the half-life of Smurf mice. According to the Nyquist–Shannon sampling theorem^35^, sampling must occur at least twice the signal frequency to avoid information loss. In flies, we assessed Smurf status every 24 hours, for a Smurf life expectancy of about 2.5 days, which allowed us to detect 100%^1,5^ of Smurf individuals (50% in Bitner et al.^4^). Beyond sampling frequency, although we chose the FITC-Dextran gavage test based on conclusions identifying it as the most suitable method for repeated in vivo permeability measurements^26^, it may not have been the most appropriate method for testing our specific hypothesis. Unlike the continuous administration of blue dye in Drosophila, which allows for accumulation in the hemolymph and thus reliable detection, the FITC test is administered as a single acute dose and does not permit this accumulation. In addition, our data analysis strategy aimed at developing a model that could generalize Smurf identification across both tested genetic backgrounds and sexes: we therefore chose to retain only six physiological parameters that exhibited consistent behavior across all conditions. While this approach aimed to ensure robustness and comparability, it may have resulted in the loss of potentially informative variables. As a consequence, this reduction in parameter diversity could lead to decreased sensitivity in detecting Smurf individuals.

Our results support the broader applicability of the two-phase ageing model. By demonstrating the existence of a Smurf state in mice, we show that the two-phase ageing model - originally developed in flies and later extended to nematodes and zebrafish - can also be applied to a mammalian system. This opens the possibility to reanalyze previously published studies to search for changes associated with this late-life phase and to identify other potential correlates - such as circulating DNA, hormones, metabolites, etc. For example, an analysis we performed using glycemia data from mice, originally published by Palliyaguru et al.^31^, revealed two distinct phases in the aging process, with a decline in glycemia preceding death, as illustrated in Supplementary Figure 5 of our study. This finding supports the robustness of our model, as it emerged without any prior hypothesis from physiological data collected at two independent sites. It also reinforces the idea that relatively small cohorts may still be sufficient to detect and capture key aspects of the ageing trajectory.

These observations, together with previously published datasets^31^ supporting a similar two-phase ageing pattern, raise the question of how our approach compares to existing ageing models and markers. Over the past decade, many ageing markers have been developed - including molecular predictors of mortality risk^20,22^, frailty indices^36^, and, more recently, so-called “ageing clocks ^24^”. These tools are often grouped under the broadly accepted “hallmarks of ageing^3^”, which are defined as (1) features associated with age, (2) features whose experimental intensification accelerates ageing, and (3) features that represent potential therapeutic targets to slow, halt, or reverse ageing. While widely used in both model organisms and humans, these markers remain primarily correlative and offer limited insight into the causal mechanisms of ageing. In this context, our work adds a complementary perspective by linking physiological deterioration to a discrete and stereotyped transition. As such, this model allows an objective measurement of frailty.

In *D. melanogaster*, we recently showed that transcriptomic changes in Smurf individuals follow a reproducible pattern, which allows us to separate time-dependent effects from physiological ageing in transcriptional hallmarks^6^. Most molecular changes, previously described as age-related^3,37^, appear instead to be Smurf-related, challenging current interpretations of the molecular basis of ageing. Extending this ageing model to mammals will help disentangle age-related changes, due to chronological age, from those characteristic due to physiological ageing. This is exemplified by our gut microbiota analysis in AKR/J mice, stratified by age (6-7 months, end of the mortality plateau; 9-12 months, T50) and Smurf status. This approach revealed distinct and unexpected microbial changes. Bacterial families such as *Bifidobacteriaceae*, *Eubacteriaceae*, and *Desulfovibrionaceae* showed increased abundance with age in non-Smurf individuals, an effect absent in Smurfs, suggesting a time-related signature (Supplementary Figures A-C). In contrast, families like *Bacteroidaceae*, *Oscillospiraceae*, *Eggerthellaceae*, and *Eubacteriaceae incerta sedis* remained stable with age but shifted specifically in Smurf individuals, indicating a profile associated with late-life physiological decline (Supplementary Figures D-G). Lastly, Lactobacillaceae showed a decrease between 6-7 and 9-12 months driven solely by Smurf mice, with no change in non-Smurfs, further supporting a signature of terminal ageing rather than chronological time (Supplementary Figure H). Aging in humans and mice is associated with complex shifts in the gut microbiome composition at the family level. Several studies in aging mice and humans have consistently reported enrichment of potentially harmful bacterial families such as Desulfovibrionaceae with advancing age, accompanied by a decline in beneficial genera such as *Lactobacillus*^39–41^. The *Lactobacillaceae* family, which we identified in our study as a biomarker of late-stage aging, has previously been linked to deteriorating health and increased frailty in the elderly, suggesting that specific species within this family may play a significant role in health status during aging^38^. Altogether, these findings suggest that not all age-related changes of the microbiome are strictly time-dependent, and that physiological ageing can reveal distinct biological signatures that remain hidden when considering chronological age alone.

Our study raises the question of whether this process is conserved in humans, and what the broader societal implications might be. Increased intestinal permeability has already been observed in terminally ill^16^ and septic patients^15^, but our model suggests that this may be more than just a symptom of terminal illness—it may serve as a valuable tool for understanding broadly conserved ageing mechanisms. While we do not claim that our model is directly generalizable to humans, our findings support the rationale for its further exploration. In this view, we recently started the clinical trial BlueRea (ClinicalTrialID NCT06845865) aiming at validating the model to predict the outcome for patients of intensive care units.

## Supplementary figures

**Supplementary Figure 1:**
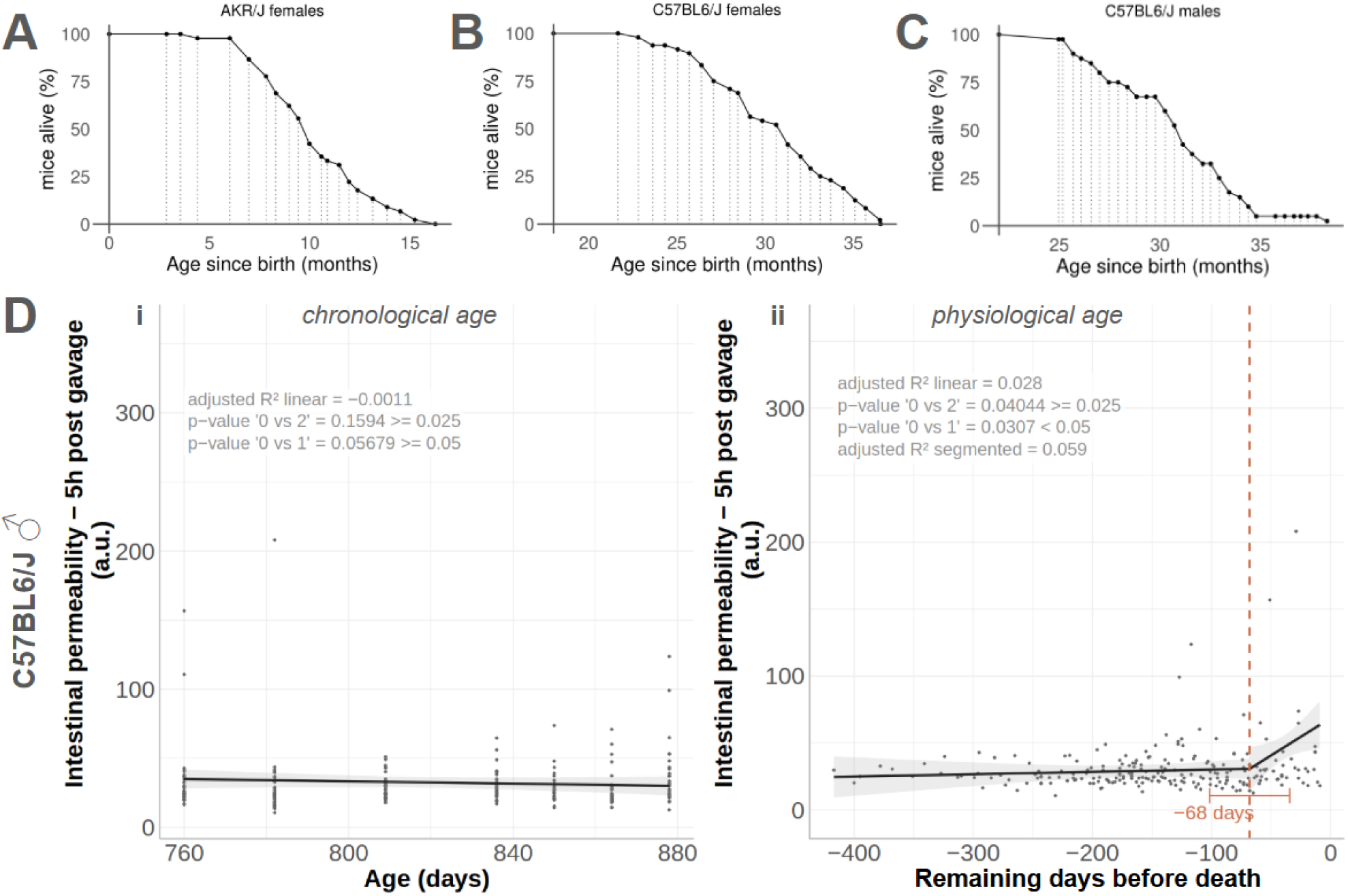
Segmented linear model analysis of physiological variables as a function of remaining days before natural death of individuals in C57BL:6J male mice cohort. Variables are: glycemia (a), body weight (b), fat mass expressed as a percentage of body weight (c), average daily energy balance (d, food intake - energy expenditure), food intake (e), locomotor activity (f), energy expenditure (g) and rectal temperature (h).Survival curve of AKR/J females cohort (n=45), (B) C57BL6/J females cohort (n=48) and (C) C57BL6/J males cohort (n=40). Dashed vertical grey arrows intersecting the survival curve of the cohort represent the time points where physiological parameters were monitored. Fluorescence unit in plasma 5h after FITC-dextran gavage indicating intestinal permeability in C57BL6/J male cohort (n=40) as a function of chronological age of individuals (i) and as a function of physiological age (ii) meaning days remaining before natural death of individuals. The dataset was modified to alleviate the impact of inter-individual variability by negating the intercept, slope, and slope differences. This treatment does not affect the pre-death pattern of the data. The vertical dashed orange line shows the breakpoint identified by segmented linear model (SLM)^42^ analysis of longitudinal intestinal permeability values. Black lines represent the results of SLM analysis before and after the breakpoint. Results of subsequent calculations and statistical tests (p-values) are written on the graph : adjusted R^2 for the linear and segmented fit ; statistical test for the absence of 1 breakpoint (BP)^28^ and for the absence of a 2nd BP (Pseudo-score test).

**Supplementary Figure 2:**
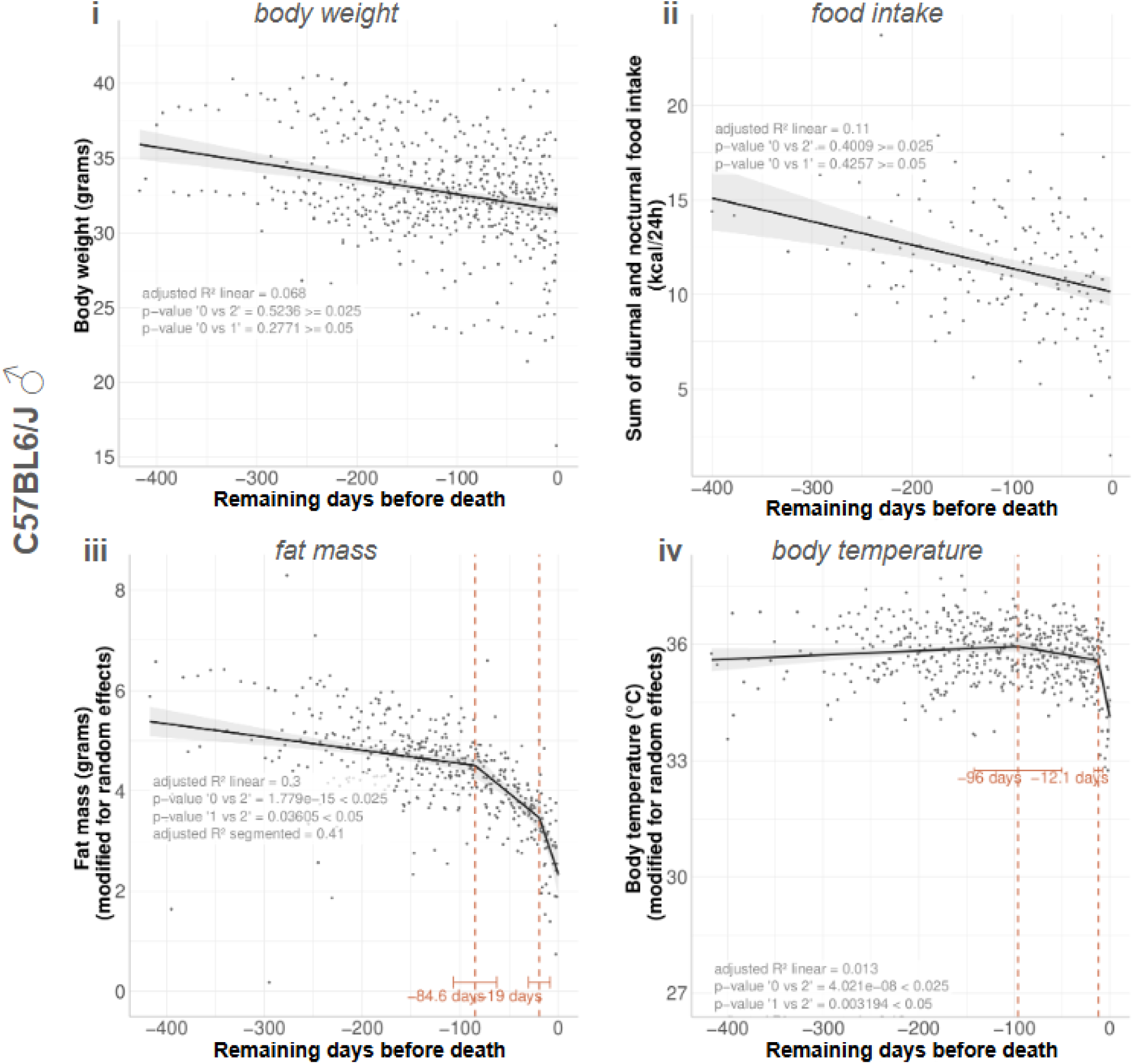
Segmented linear model analysis of physiological variables in a male C57BL/6J mice cohort. Longitudinal representation of body weight (i), food intake (ii), fat mass (iii) and rectal temperature (iv) all expressed as a function of time before recorded death (n=40). The dataset was modified to alleviate the impact of inter-individual variability by negating the intercept, slope, and slope differences. This treatment does not affect the pre-death pattern of the data. The vertical dashed orange line shows the breakpoint identified by segmented linear model (SLM) analysis of longitudinal intestinal permeability values. Black lines represent the results of SLM analysis before and after the breakpoint. Results of subsequent calculations and statistical tests (p-values) are written on the graph : adjusted R^2 for the linear and segmented fit ; statistical test for the absence of 1 breakpoint (BP)^28^ and for the absence of a 2nd BP (Pseudo-score test).

**Supplementary Figure 3:**
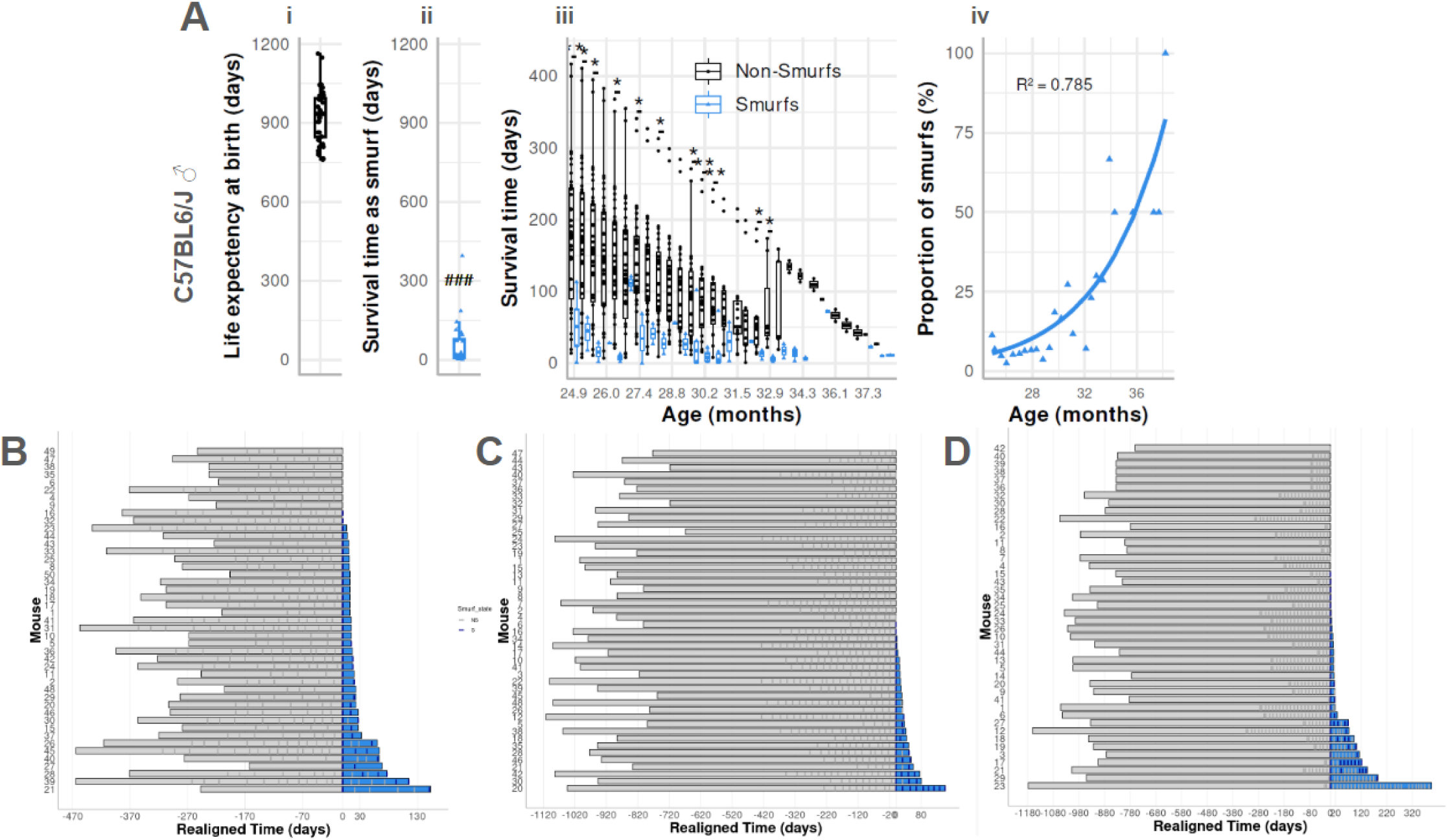
Identifying individuals with the same chronological age but different mortality risks in C57BL/6J male mice (n=40). (Ai) Individual life expectancy at birth in days (black boxes, points); (Aii) Individual survival time as smurf in days (blue boxes, triangles); (Aiii) Individual survival time as non-smurf (black boxes, dots) and as smurf (blue boxes, triangles) as a function of chronological age. The box plot represents the interquartile range, showing the middle 50% of the data from the first to the third quartile. The line inside the box marks the median, and whiskers extend to the smallest and largest values. Statistical significance: ### p < 0.001 by paired Wilcoxon test comparing mean life expectancy at birth and mean survival time as smurf (ii); *p < 0.05, **p < 0.01, ***p < 0.001 by Wilcoxon test comparing mean survival times of non-smurf and smurf individuals (iii) (Aiv) Proportion of smurf individuals as a percentage of the living population (blue triangles), with an exponential growth regression of smurf proportion (blue line). (B, C, D) Representation of individual mice life histories followed from adulthood to their natural death, as,and centered on the moment of the smurf transition in AKRJ females (B), C57BL6/J females (C) and C57BL6/J males (D). Grey rectangles indicate mice life before smurf transition and blue rectangles display mice life after the smurf transition. . In the AKR/J female cohort, 7 individuals were never classified as smurfs, and 2 out of 38 were scored non-smurf after their initial categorization as smurf. In the C57BL6/J female cohort, 24 individuals were never classified as smurfs, and 5 out of 24 were scored non-smurf after their initial categorization as smurf. In the C57BL6/J male cohort 16 individuals were never classified as smurfs, and 12 out of 24 were scored non-smurf state their initial categorization as smurf. SLM = Segmented Linear Model; PCA = Principal Component Analysis; Dim1 = Dimension 1 of the PCA ; R2 = 1 - (sum squared regression / total sum of squared)

**Supplementary Figure 4 :**
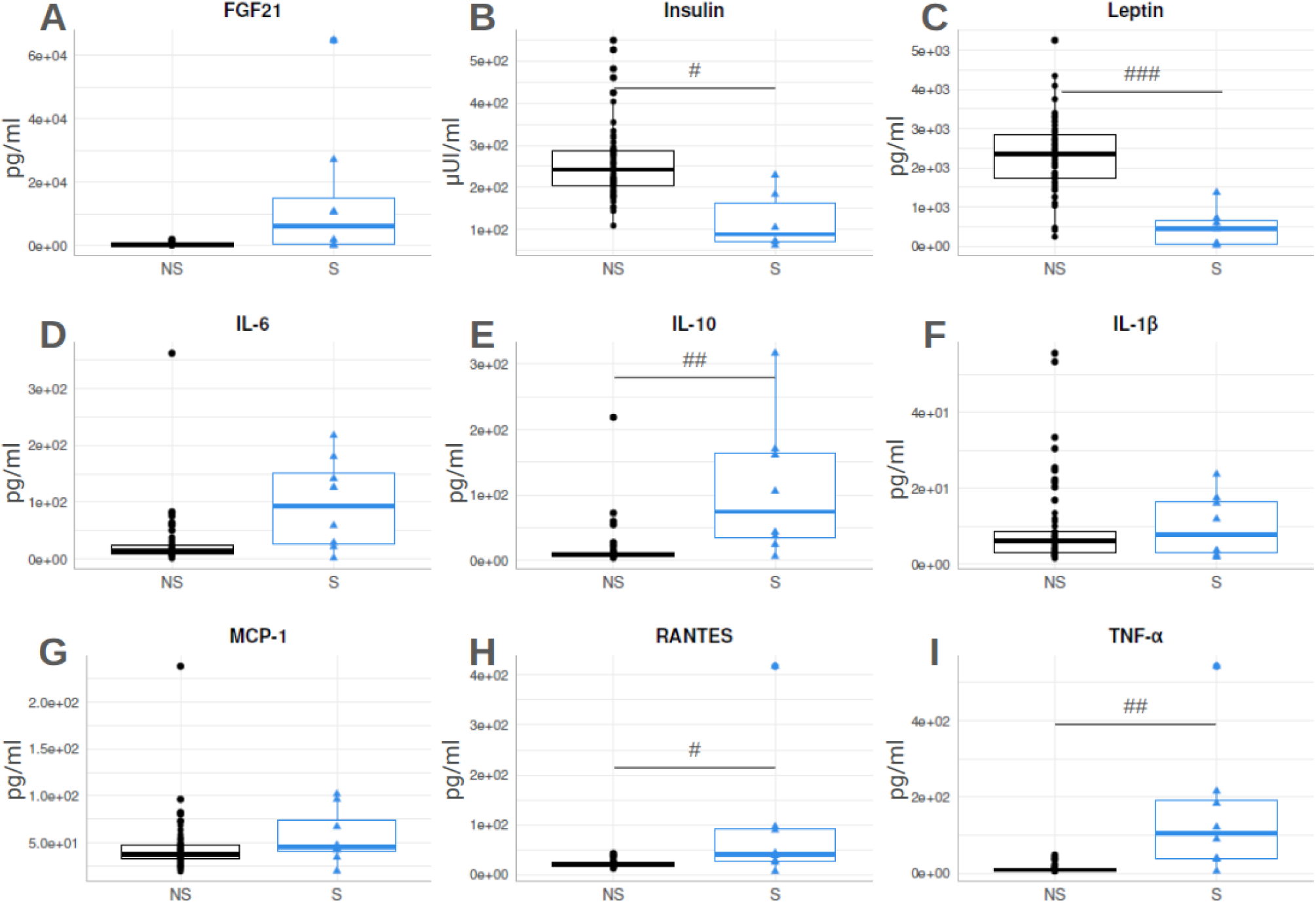
Plasma concentrations of hormones, cytokines and chemokines in the AKR/J female mouse cohort. Fibroblast growth factor 21 (A), insulin (B), leptin (C), interleukin 6 (D) interleukin 10 (E), interleukin-1 beta (F), monocyte chemoattractant protein-1 (MCP-1/CCL2) (G), Regulated on Activation, Normal T cell Expressed and Secreted (RANTES/CCL5) (H), Tumor Necrosis Factor alpha (I). Results are shown according to individual status: NS (non-smurf) or S (smurf), as defined in the main text. The boxes represent the interquartile range, spanning the first to third quartiles, with the horizontal line indicating the median. Whiskers extend to the minimum and maximum observed values. Individual data points are overlaid to show the variability within each group. Statistical comparisons of mean plasma concentrations between groups were performed using an unpaired Wilcoxon test, with p-values adjusted for multiple testing using the Bonferroni correction. Statistical significance is denoted as follows: # p < 0.05, ## p < 0.01, ### p < 0.001. Group sizes: NS (n = 60), S (n = 8).

**Supplementary Figure 5:**
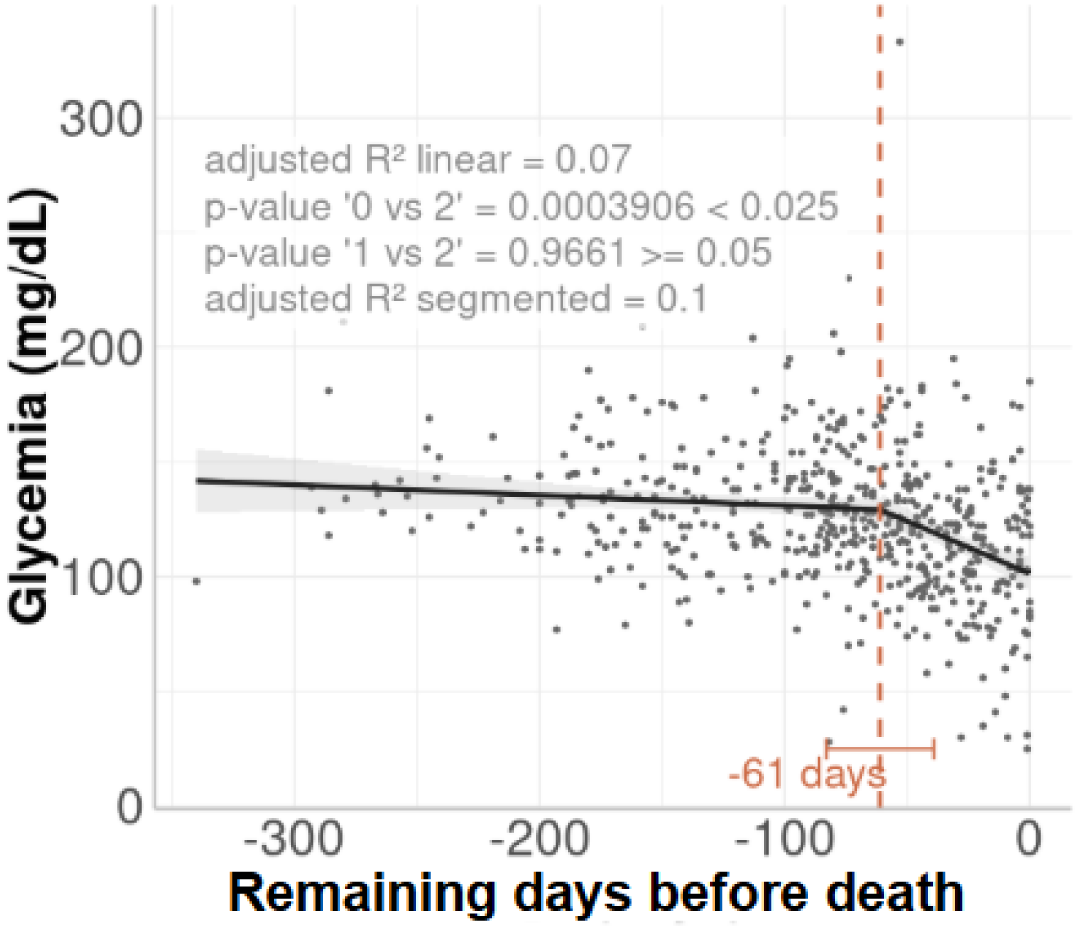
Segmented linear model analysis of plasmatic glucose concentration data from Palliyaguru et al. (PMID34508697) as a function of remaining days before natural death of individuals. Variable is glycemia. The dataset is obtained by independent measurements on individual mice. The vertical dashed orange line shows the breakpoint identified by segmented linear model (SLM) analysis of longitudinal intestinal permeability values. Black lines represent the results of SLM analysis before and after the breakpoint. Results of subsequent calculations and statistical tests (p-values) are written on the graph : adjusted R^2 for the linear and segmented fit ; statistical test for the absence of 1 breakpoint (BP)^28^ and for the absence of a 2nd BP (Pseudo-score test). n=530 HET3 & B6 - female (n=238) male (n=292) mice. Data from: Fasting Blood Glucose as a Predictor of Mortality: Lost in Translation^43^.

**Supplementary Figure 6:**
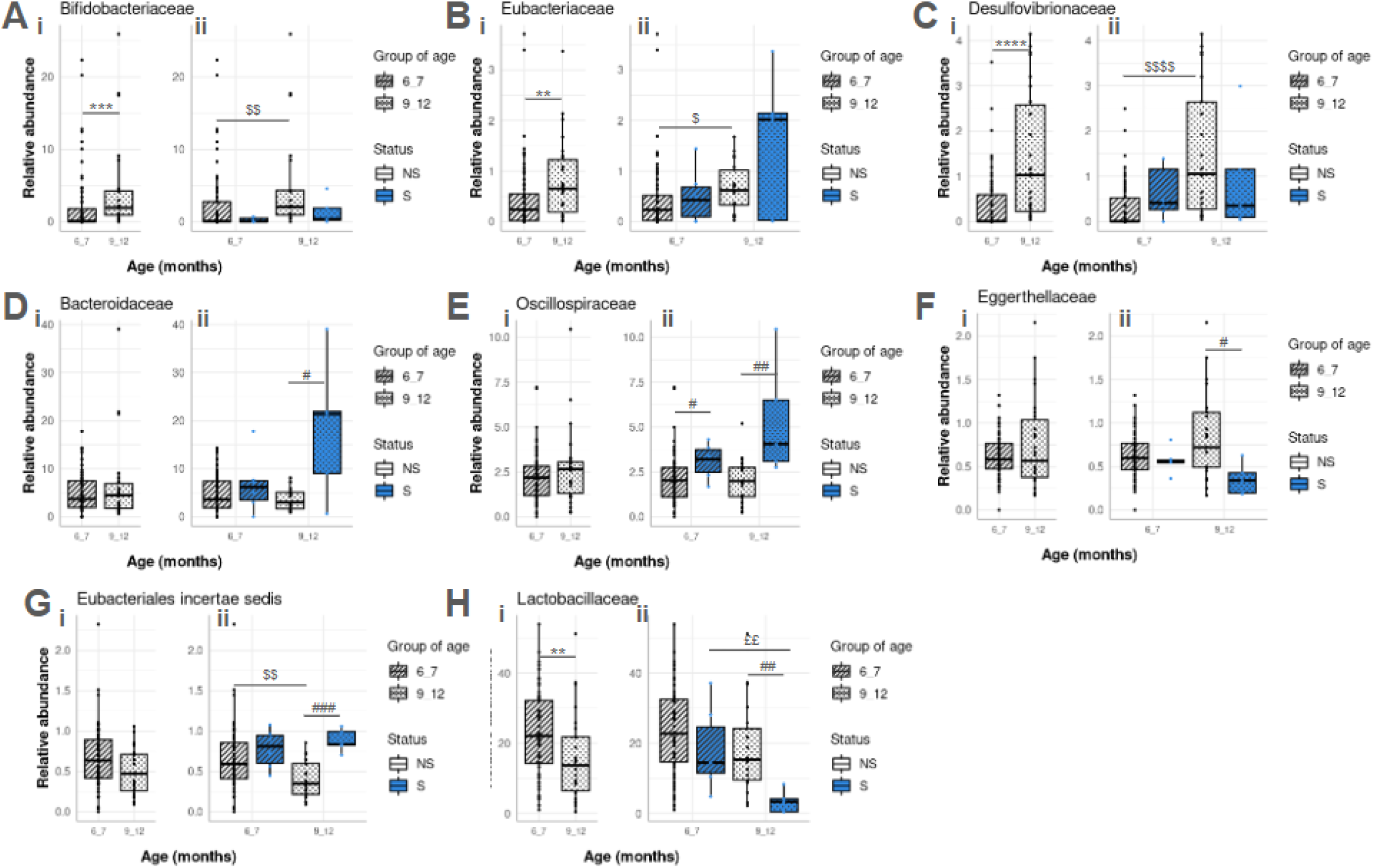
Gut microbiota family-level relative abundances in the AKR/J female mouse cohort. Results are presented either based solely on the chronological age of individuals, grouped into 6–7 or 9–12 months, or further stratified within each age group according to status — NS (non-smurf) or S (smurf) — as defined in the article. The box plot represents the interquartile range, showing the middle 50% of the data from the first to the third quartile. The line inside the box marks the median, and whiskers extend to the smallest and largest values. Comparisons of mean relative abundances were performed using the unpaired Wilcoxon test (no p values adjustment). Statistical significance is indicated as follows: * p < 0.01, ***p < 0.001, ****p < 1e-04 for comparisons between age groups “6_7” and “9_12”; #p < 0.05, ##p < 0.01, ###p < 0.001 for comparisons between NS and S status across all age groups; $p < 0.05, $$p < 0.01, $$$$p < 1e-04 for comparisons between age groups “6_7” and “9_12” within the NS group; and ££p < 0.01 for the same comparison within the S group. Group sizes – NS 6–7 months : n=58, NS 9–12 months : n=21, S 6–7 months : n=6, S 9–12 months : n=5.

## Materials and Methods

### Protocol

#### Animals

Fifty 10-week-old AKR/J female mice (Charles Rivers, US), fifty 20–24-month-old C57BL/6J male mice and fifty 20–24-month-old C57BL/6J female mice (Janvier Labs, France) were housed in groups (five animals per cage) or individually in PET plastic cages (EasyCage® mouse, Allentown Inc, France) with corn cob bedding (Lab cob 12, Serlab, France) at 21 ± 1°C and 55 ± 5% humidity, under a 12-hour light / 12-hour dark cycle. The mice acclimated for 2–3 weeks before experiments. Food and water was provided ad libitum (3.339 Kcal/g with 19.3% from proteins, 8.4% from lipids, and 72.4% from carbohydrates; SAFE#A04) and animals were monitored daily. According to the project’s ethical approval, we euthanized any animal that (1) showed clinical signs of pain, (2) developed an ulcerated or disabling tumor, or (3) had a tumor exceeding 1.2 cm in diameter, using neck dislocation under isoflurane anesthesia. AKR/J females were group-housed throughout the experiment, except during metabolic measurements (PhenoMaster - TSE System), when we had to individualize them temporarily. After working with this first group of AKRJ females, we wanted to reduce the slight weight loss caused by repeated individualization. Therefore, we isolated 40–50% of the C57BL/6J mice upon arrival and kept them isolated until natural death. The exact percentage depended on the number of metabolic cages available at the start of the experiment. These isolated mice entered the metabolic cages every 3–4 weeks. As mice from the isolated group died, we isolated additional animals from the group-housed cohort to maintain enough individuals for metabolic recordings (Promethion Core - Sable System). We isolated these mice at least one week in advance to limit the effects of isolation on physiological parameters. For all C57BL/6J males and females, we recorded body weight and composition, glycemia, plasma and fecal samples, intestinal permeability, and rectal temperature every 2–3 weeks from arrival until natural death. We documented whether each measurement came from a grouped or isolated mouse in the dataset. For ethical reasons, we euthanized about 30% of the animals. Based on these observations, we took several actions: (1) we decided not to repeat the experiment with a male AKRJ group as originally planned and continued using only C57BL/6J mice, (2) we reclassified our ethical protocol as severe, and (3) we kept data from euthanized animals to analyze physiological changes before death, considering that death would have occurred within 24 hours.

#### 0.8 kDa FITC-Dextran synthesis

Glucosamine hydrochloride (5.4 mg, 0.025 mmol) and fluorescein isothiocyanate (FITC, 10 mg, 0.025 mmol) were dissolved in DMF (1mL). DIPEA (9 μL, 0.05 mmol) was added and the mixture was stirred at 40°C for 24 hours. The mixture was added to DCM dropwise and the precipitate was collected by filtration and crystallized on MeOH/Cyclohexane 1/1 to give the product as a yellow/red solid (8.8 g, 62%). 1H NMR (400 MHz, D2O) δ 8.16 (d, J = 1.9 Hz, 1H), 7.93 (dd, J = 8.1, 1.9 Hz, 1H), 7.27 (d, J = 8.1 Hz, 1H), 6.79 (dd, J = 8.1 Hz, 2.3Hz, 1H), 6.77 – 6.69 (m, 3H), 6.65 – 6.60 (m, 2H), 5.71 (d, J = 2.2 Hz, 1H), 4.87 (d, J = 6.0 Hz, 1H), 4.53 (d, J = 7.8 Hz, 1H), 4.52 (d, J = 6.0 Hz, 1H), 4.05 (m, 2H), 3.9-3.8 (dd, J =3.3-10.8 Hz, 2H), 3.78-3.65 (m,4H).

#### FITC-Dextran solution preparation

At the time of experiment the smallest size of FITC-Dextran on the market was 4kDa. For the experiments performed on the AKRJ group we used the solution A prepared from a mix of FITC-Dextran of different sizes: 4kDa, 20 kDa and 70 kDa. Since we did not observe a massive increase in intestinal permeability, we redesigned our protocol and collaborated with a chemist to make a FITC with a size close to the one of the Blue#1 (792.58 Da) used in the Drosophila protocol. This new FITC of 568 Da was used to make the solution B in the experiments performed on the C57BL/6J mouse lines. Solution A : fluorescein isothiocyanate–dextran powder (FITC-Dextran, TdB labs., Sweden) 4kDa, 20kDa and 70kDa (60mg/mL) were diluted together in sterile water. Solution B : FITC-Dextran 0.8 Da (See “0.8 Da FITC-Dextran production” section) was kept at a concentration of 280mg/mL in DMSO (Dimethyl sulfoxide BioUltra, for molecular biology, ≥99.5%, Sigma-Aldrich®, US) at -80°C and diluted (7.25mg/mL) in sterile water for the gavage. Both solutions were prepared the same day of the gavage in a 15 mL Falcon^TM^ tube (Corning, US), just before the test and are protected from light using foil.

#### FITC-Dextran gavage test

All mice were food-deprived for 3 hours at the beginning of the light cycle. Home bedding and water bottles were kept in the cage. When the 3 hours of food deprivation were up, body weight and glycaemia were measured, and 75 µL of blood were collected in awake animals at the tail tip after incision. Once the pre-test measurements and samples had been taken, the test started with the gavage of the mice with the FITC-Dextran solution A or B (100 µL/10 g of body mass) in awakened animals at the tail tip. Glycaemia was expressed in ng/mL and measured using a glucometer (Glucofix® Sensor and Glucofix® Premium, A. Menarini diagnostics, France). Blood was collected using haematocrit capillaries coated with heparin (Hirschmann® Laborgeräteann, Germany). Blood samples were kept on ice before centrifugation in 0.5 mL Eppendorf® Safe-Lock tubes (Eppendorf®, Germany). After centrifugation (10,000 × g for 7 min at 4°C), plasma samples were collected and immediately frozen at −80°C and kept for further analyses. Mice were left in their home cages. If they were grouped or isolated at the beginning of the test, they remained so during the test. At the end of the test, after the last sampling, food was put back in the cage of the mice. One, three, and five hours after the gavage, 20 µL blood samples were collected.

#### Fluorescent Plasma Measurements

Plasma samples were diluted 5:95 (v/v) in DPBS (Dulbecco’s Phosphate Buffered Saline, Gibco^TM^, ThermoFisher, US). Fluorescence was measured spectrophotometrically (FlexStation® 3, Molecular Devices, US) in 96-well dark plates (Non-Treated Surface, Non-Sterile, Thermo Scientific^TM^, US) using the following parameters: excitation = 493 / emission = 525 / cut off = 515 / room temperature. Fluorescence is expressed in arbitrary units.

#### Indirect calorimetry measurements

Oxygen consumption (VO₂, mL/15 min), carbon dioxide production (VCO₂, mL/15 min), food intake (g/15 min), water intake (mL/15 min), and locomotor activity (counts/h) were measured using indirect calorimetry apparatus (PhenoMaster, TSE Systems GmbH, Germany; and Promethion™, Sable Systems International, US). Mice were individually housed with ad libitum access to food and water. Measurements included a minimum of 3 up to 5 days of data recording. The first day and the first night of recording were not taken into account and were considered as a habituation period to the indirect calorimetry apparatus. We summed the measurements of food and water intake, VO₂, VCO₂, and infrared beam crossings on the X, Y, and Z axes taken every 15 minutes to evaluate hourly and daily values. This article focuses on food intake data, while other recorded parameters are accessible in the accompanying raw data

#### Body temperature measurements

The mouse was hand-restrained and a temperature probe (Physitemp thermal TH5, Phymep, France) covered with Glycerin (DUREX® Gel lubrifiant naturel, France) was gently inserted into the rectum to a fixed depth: 2 cm.The body temperature was always taken at the same time of day, between 10 and 11 am.

#### Body composition analysis

Body mass composition, lean mass, fat mass, free water and total water content are analyzed by magnetic resonance imaging with the EchoMRI 100 system or Minispec LF90 (Whole Body Composition Analyzers, EchoMRI, Bruker, US) according to the manufacturer’s instructions in awake mice.

#### Metabolic biomarkers, cytokines and chemokines quantification in plasma

A U-Plex multiplex assay (K152ACL-2, Metabolic Group1 (ms) assay, Meso Scale Diagnostics, US) was used to measure the levels of metabolic biomarkers, cytokines and chemokines (insulin, leptin, FGF-21, IL1-beta, IL-6, IL-10, MCP-1, RANTES, TNF-alpha) in mice plasma according to the manufacturer’s protocol.

#### Microbiota analysis

Fecal samples were directly collected at the exit of the anus of the hand-restrained mouse in 0.5 mL Eppendorf® Safe-Lock tubes (Eppendorf®, Germany), which were instantly placed in liquid nitrogen and kept at −80°C for further analyses. Fecal samples were always collected at the same time of day, between 10 and 11 a.m. We extracted total microbial DNA from approximately 40 mg of fecal sample using a kit for the isolation of genomic DNA from stool samples according to the manufacturer’s protocol (15573626, Macherey-Nagel™ NucleoSpin™ DNA Stool, Fisher Scientific, France). We assessed DNA concentration and integrity by spectrophotometry using the Qubit dsDNA HS (High Sensitivity) assay kit (Q32851, Qubit™ dsDNA HS, Invitrogen™, US). We amplified the V3–V4 hypervariable region of the bacterial 16S rDNA by PCR using the following primers: forward primer 5’-CTTTCCCTACACGACGCTCTTCCGATCTACGGRAGGCAGCAG-3’ and reverse primer 5’-GGAGTTCAGACGTGTGCTCTTCCGATCTTACCAGGGTATCTAATCCT-3’. Amplifications were carried out using the following ramping profile: 1 cycle at 94°C for 1 minute, followed by 30 cycles at 94°C for 1 minute, 65°C for 1 minute, and 72°C for 1 minute, ending with a final step at 72°C for 10 minutes. After quality checking by electrophoretic migration on a 2% agarose gel, we sequenced the resulting amplicons using Illumina MiSeq technology (GenoToul platform, Toulouse, France). The resulting sequences were analysed using command line with FROGS v5.0.0^44^ on the Migale Bioinformatics server (Université Paris-Saclay, INRAE, MaIAGE, 78350, Jouy-en-Josas, France). Reads were filtered and merged to generate ASVs with dada2.

The guidelines of FROGS v5.0.0 were followed to detect and remove chimeras using the vsearch tool. ASVs with global abundance lower than 0.005 % were removed from the following analysis with FROGS filters tool and the amplicon sequence variants (ASV) in the sequence table were then assigned to species using FROGS affiliation with NCBI 16S Ribosomal RNA REFseq Bacteria (v1.20230726) database. Microbiota bioinformatic analysis was performed on R v4.3.3 with package Phyloseq v1.46.0, ggplot2v3.5.1 and phyloseq.extended v0.1.4.1. We used raw, un-rarefied ASVcounts to identify taxa abundances and characterize gut microbiota composition. Family relative abundances were compared using a Mann-Whitney test.

### Sampling plan

#### Power calculation

For this *a priori* power analysis, we considered the case of detection of a mean difference between the "Smurf" and the "non-Smurf" groups at T_50_, when 50% of the population is dead, assuming 30% of “Smurf” individuals are alive based on drosophila studies and our mathematical model^5^. A total sample size N = 50 would correspond to 17 "non-Smurf" and 8 "Smurf" individuals at T_50_. An effect size of 1.6 can be detected with ɑ = 0.05 and 1-*β* = 0.95, (G*Power 3.1.9.731, Sensitivity analysis) at 80% POWER. This is considered an extra-large effect according to Cohen’s criteria^45^, but is consistent with results obtained in Drosophila^5^. Thus, our proposed sample size of N=50 was adequate for this study, and was approved by the Animal Care Committee (see Ethical Approval Plan for details). G*Power 3.1.9.7 uses the equations described in Cohen, J. Statistical Power Analysis for the Behavioral Sciences (Academic Press, 2013)^46^.

#### Criteria for data exclusion

Regarding body weight and composition, glycemia, body temperature, plasmatic fluorescent level, as well as metabolic biomarkers, cytokine and chemokine quantification in plasma, and microbiota composition, we excluded no data. When possible, at the time of measurement, if a value fell outside its expected range, we immediately performed a second measurement to validate the first. If the second measurement did not match the first, we performed a third measurement and considered the average of the two closest values. In very rare cases, when blood glucose levels dropped below 20 mg/dL, the glucometer could not display the value and showed “LOW” on the screen. In such cases only, we assumed the glucose value to be 20 mg/dL. For O₂ and CO₂ measurements, locomotor activity, food, and water intake, the simultaneous analysis of metabolic and behavioral data helped us detect inconsistencies and errors, thus preventing the inclusion of false data. In such cases, we did not replace the data but removed them from the study. Accidental death of an animal due to an external factor leads to its exclusion from analyses based on the number of days remaining before death, but not from analyses based on chronological age.

If an animal near death presented with a deteriorated clinical condition that required ethical euthanasia, we retained its full dataset for certain analyses. Finally, fitting the segmented linear mixed effect model (SLME) required at least four data points per mouse; otherwise, the model became misspecified and yielded no solution. Consequently, for each outcome variable, we removed all mice with three or fewer data points for that variable. In all cases, we specified this in the corresponding figure legend.

#### Cessation of data collection

Data collection stopped when all individuals in each group have died naturally or been euthanized for ethical reasons defined in APAFIS#18333-2018112915281820v6.

### Analysis pipeline

#### Distinguish the two phases 1 and 2 of ageing in the mouse

In essence, we fit a segmented linear regression model (SLM) for each biomarker of interest in order to identify a two-phase evolution. The model can be fitted with zero, one, or multiple breakpoints (BPs). A two-phase evolution is supported if the model with one breakpoint provides a good fit (*R*^2^_segmented_>*R*^2^_linear_) i.e. if the slopes before and after the breakpoint are estimated to be significantly different (p-value testing the null hypothesis of a BP > 0.05). One specific issue here is the longitudinal nature of the data, each mouse being measured several times. Hence we use the following procedure: (1) Fit a segmented linear mixed effect model (SLME) and segmented linear model (SLM) and compare the variance of breakpoint estimates; (2) If the mixed effect is significant, use the estimated mixed effects to compute a modified explanatory variable where the mixed effect of each mouse has been zeroed-out (intercept, slope, and slope difference), else, go directly to the next step ; (3) Fit an SLM and statistically test the non-nullity of the slope difference at the breakpoint (i.e. the presence of 1 breakpoint versus 0 breakpoint) and that there is only one breakpoint (i.e. the presence of 1 breakpoint versus 2 breakpoints). These analyses allow us to test the existence of two phases during ageing with a specific physiologic signature of the end of life in the mouse model. Change in data behavior before and after the breakpoint can also be tested using a repeated measures analysis with all measurements by splitting data into two groups: "phase 1" group before the breakpoint and "phase 2" group after the breakpoint, and treating mice as a random variable to take into account native differences among mice, and including an autocorrelation structure. We will add the variable “single/group housed” as a covariate in both the SLM and SLME models. This will allow us to test for the presence of a breakpoint and to estimate its position while taking into account the potential effect of single/grouped housing. SLMs are fitted using the R package “segmented”. For better visualization and to graphically illustrate the trend in our data points (not to infer any model parameter) figures may also include a nonparametric local regression estimate of each variable, using locally estimated scatterplot smoothing (LOWESS^47^).

#### Differentiate individuals with the same chronological age but with different life expectancy

If one or more breakpoints were identified, we calculated an average breakpoint across the different physiological variables. We selected physiological markers showing a breakpoint to perform a principal component analysis (PCA). We used PCA results to determine whether individuals were in phase 1 or 2 of ageing at any given chronological age, based on the score of the dimension 1. After fitting a segmented linear model (SLM) on the score of the dimension of interest, we calculated the PCA threshold value as follows: PCA_threshold = (Mean of Dimension Score in phase 1) +/- (1.65 × Standard Deviation of Mean in phase 1). Depending on data behavior between phases 1 and 2 of ageing, we defined individuals with a score below or above this PCA_threshold value as “non-Smurf” if they were in phase 1, and “Smurf” if they were in phase 2 of ageing at any given chronological age.

### Statistical analysis

We use the R package “segmented” to fit an SLM and an SLME^27^. The graphical representation of the model residuals displays an (approximately) normal distribution. The implementation of the SLM comes with a statistical test for the presence of one breakpoint versus zero breakpoint (pseudo score statistical test^28^); selection criteria for the correct number of breakpoints (Akaike Information Criterion (AIC)^48^); a statistical test for the nullity of the difference of slope at the breakpoint (Davie’s test^28,49^). However, SLME provide no such tests. So, in order to test for the presence of a breakpoint while taking mixed effects into account, we use the estimated mixed effects to compute a modified explanatory variable where the mixed effect of each mouse has been zeroed-out. This modified dataset presents the same pattern as the data, except that the effect of each mouse of the intercept, slope, and slope difference has been negated. Additionally, we can compute the adjusted *R*^2^ coefficient for the segmented fit and compare it to that of the LM to assess the benefit of using the latter model. The width of the 95% confidence interval around the pointwise estimate of the breakpoint also helps to assess the confidence in the model fit.

It is worth noting that from a statistical standpoint, we use the modified data as if it were the real data, without taking into account the uncertainty stemming from the estimation of the mixed effect. This boils down to slightly underestimating the uncertainty of the estimation provided by the SLM. We reckon that this has minimal practical implications as the p-values obtained for the non-nullity of the slope difference are usually very small.

We use the “FactoMineR” R package for PCA analysis. Comparison between “non-Smurf” and “Smurf” groups were carried out using the Wilcoxon test (unpaired). In Supplementary Figure 6, comparing Smurf and non-Smurf individuals was challenging due to partially paired samples. As some individuals were measured multiple times and others not at all, we used an unpaired Wilcoxon test to minimize the risk of false positives. All tests were performed using R R6_2.6.1, Microsoft Office Excel 2016, G*Power 3.1.9.7 and JASP 0.15.0.0^50^. Numbers of animals are given in the legends. All scripts are available online.

The measurement tools we use ensure that we cover the whole range of values that each longitudinal variable can adopt. This guarantees the absence of floor or ceiling effects in their distributions. The only sensitive parameter is the measurement of blood glucose by the glucometer, which does not allow measurements below 20ng/ml. However, this is not a problem because a blood sugar level below this value is very rare and indicates imminent death, which will not bias our interpretations.

#### Joint model

To assess whether biomarkers identified at stage 1 as showing a breakpoint independently predicted impending death, we adjusted joint models^51^ using the R package JMBayes2. Briefly, a joint model simultaneously models the longitudinal evolution of biomarkers of interest and time-to-death, allowing for the investigation of how changes in biomarker profiles modify the risk of death. Included biomarkers were body temperature, fat content, glycemia, and 3-5 hour intestinal permeability. Models were adjusted separately for C57 and AKRJ mouse. For C57, we adjusted on sex in both the longitudinal and survival models. For all biomarkers, we accounted for non-linearities in the longitudinal profile using natural cubic splines of chronological age.

## Data Availability Plan and Conflict of Interest

All data and codes are available on GitHub https://github.com/MichaelRera/BMCBiol_SmurfMice.git. This applies to all data and other information about mice (IDs, data on single-versus group-housed status, physiological and metabolic data…).The authors declare no conflict of interest.

## Ethical Approval Plan

All protocols are carried out in accordance with French standard ethical guidelines for laboratory animals and with approval of the Animal Care Committee (Comité d’éthique en expérimentation animale Buffon C2EA-40; Comité d’éthique pour l’expérimentation animale Charles Darwin CEEACD/N°5 ; APAFIS#18333-2018112915281820v6 ; APAFIS#30102-2021022517524316v4 ; “Non-technical summaries of notified files” n°10766 (www.enseignementsup-recherche.gouv.fr))

## Contributions

Michael Rera and Céline Cansell conceived and planned the presented idea, model and wrote the manuscript. Céline Cansell ran the experiments. Fanny Bain conducted the longevity experiments. Flaminia Zane provided technical support for the analysis. Vivien Goepp built the statistical pipeline for the automated identification of breakpoints in parameter values. Nicolas Todd built the joint model for predicting risk of impending death in individual mice. Veronique Douard, Magali Monnoye, Carole Rovere and Clara Sanchez ran microbiota analysis and plasma circulating markers. Nicolas Pietrancosta synthetised the 600Da FITC-dextran. Raphael GP Denis and Serge Luquet helped with the metabolic exploration platform.

## Acknowledgement

We deeply thank Dimitris Rizopoulos for his help and responsiveness regarding joint models.

